# Genomic and antigenic diversity of carried *Klebsiella pneumoniae* isolates mirrors that of invasive isolates in Blantyre, Malawi

**DOI:** 10.1101/2021.10.07.463515

**Authors:** Joseph M. Lewis, Madalitso Mphasa, Rachel Banda, Mathew A Beale, Jane Mallewa, Eva Heinz, Nicholas R Thomson, Nicholas A Feasey

## Abstract

*Klebsiella pneumoniae* is an antimicrobial resistance (AMR) associated pathogen of global importance, and polyvalent vaccines targeting *K. pneumoniae* O-antigens are in development. Genomes from sub-Saharan Africa (sSA) are underrepresented in global sequencing efforts. We therefore carried out a genomic analysis of extended-spectrum beta-lactamase (ESBL)-producing *K. pneumoniae* complex isolates colonising adults in Blantyre, Malawi, placed these isolates in a global genomic context, and compared colonising to invasive isolates from the main public hospital in Blantyre. 203 isolates from stool and rectal swabs from adults were whole-genome sequenced and compared to a publicly available multicountry collection of 484 *K. pneumoniae* genomes sampled to cover maximum diversity of the species, 150 previously sequenced Malawian and 66 Kenyan isolates from blood or sterile sites. We inferred phylogenetic relationships and analysed the diversity of genetic loci linked to AMR, virulence, capsule (K-) and LPS O-antigen (O-types). We find that the diversity of Malawian *Klebsiella* isolates is representative of the species’ population structure, but with local success and expansion of sequence types (STs) ST14, ST15, ST340 and ST307. Siderophore and hypermucoidy genes were more frequent in invasive versus carriage isolates (present in 13% vs 1%, p < 0.001) but still generally lacking in most invasive isolates. The population structure and distribution of O-antigen types was similar in Malawian invasive and carriage isolates, with O4 being more common in Malawian isolates (14%) than in previously published studies (2-5%). We conclude that host factors, pathogen opportunity or alternate virulence loci not linked to invasive disease elsewhere are likely to be the major determinants of invasive disease in Malawi. Distinct ST and O-type distributions in Malawi highlights the need for geographically aware sampling to robustly define secular trends in *Klebsiella* diversity. Colonising and invasive isolates in Blantyre are similar and hence O-typing of colonising *Klebsiella* isolates may be a rapid and cost-effective approach to describe global diversity and guide vaccine development.

**Data Summary:** All data and code to replicate this analysis is available as the *blantyreESBL* v1.0.0 R package (https://doi.org/10.5281/zenodo.5554082) available at https://github.com/joelewis101/blantyreESBL. Reads from all isolates sequenced as part of this study have been deposited in the European Nucleotide Archive, and accession numbers (as well as accession numbers of publicly available genomes used in this analysis) are provided in the R package.

## Introduction

*Klebsiella pneumoniae* is a highly prevalent human gut colonizer^1^ and opportunistic pathogen^2^ which is often significantly associated with antimicrobial resistance (AMR) and has been identified by the World Health Organisation as a global priority AMR pathogen^3^. In Low- and Middle-Income Countries (LMIC) such as the nations of sub-Saharan Africa (sSA), AMR *K. pneumoniae* presents a significant therapeutic challenge. Malawi is a low-income country in South-East Africa, where 91% of *K. pneumoniae* infections are now resistant to third-generation cephalosporins^4^ (3GC), largely mediated through production of extended-spectrum beta-lactamases (ESBLs). In this and many other LMIC, 3GC are first-line antimicrobials for severe febrile illness and alternatives with activity against ESBL-producers are often unavailable, rendering ESBL *K. pneumoniae* infections *de facto* untreatable with locally available antimicrobials.

Whole genome sequencing (WGS) has provided significant insight into the population structure of *K. pneumoniae*, which we now understand is a species complex encompassing several subspecies^5^. Despite this diversity, WGS highlighted that the global spread of AMR is linked to clonal expansion of AMR-associated high-risk clones^2^ and genomic loci associated with virulence^6^ (including the hypermucoid phenotype^7^) have been identified. Historically, antimicrobial resistance and virulence were associated with different *K. pneumoniae* populations, but convergence of AMR genes in hypervirulent lineages is increasingly described, especially in South and South-East Asia^8^, resulting in community-acquired widely-disseminated or deep seated infections in otherwise healthy individuals that are difficult to treat^9^.

In response, *K. pneumoniae* vaccines are in development^10^, targeting LPS O-antigens, which can be predicted via a sequence-based typing scheme^11,12^. Analyses of large-scale genome collections have provided important insights into the distribution and diversity across the species complex, which is essential to focus efforts to the clinically most relevant types^11,13^. However, whilst an initial ‘global’ collection^5^ represented a milestone in *K. pneumoniae* genomics and was designed to cover the diversity in a multi-country effort providing our first insight into the genomic plasticity, it is restricted to isolates from 12 countries, notably lacking any sSA representatives. Follow-up studies over the past years have also often focused on HIC clinical studies^14,15^, and genomes from sSA are drastically under-represented in genome datasets. There is an urgent need to investigate the genomic epidemiology of *K. pneumoniae* in this setting to assess whether conclusions from largely HIC collections are valid for LMICs, a crucial requirement for a vaccine to be effective in these settings where, arguably, it may have the most benefit.

In addition, though colonisation with *K. pneumoniae* is thought to precede infection in many cases^1^, sequencing efforts have largely focused on invasive isolates. There is some evidence from elsewhere in the world that colonising and invasive isolates differ^2^; understanding this difference in sSA could help to define the determinants of infection in this setting. We therefore present the results of a genomic analysis of *K. pneumoniae* from a study of colonisation with ESBL Enterobacterales in Blantyre, Malawi, with three aims: to describe the population structure, serotype diversity, AMR and virulence determinants of colonising *K. pneumoniae* in this setting; to compare colonising to previously sequenced Malawian invasive isolates; and to relate these data to observations made in other parts of the world.

## Methods

The isolates analysed in this study were colonising isolates selectively cultured from stool and/or rectal swabs collected from adults in Blantyre, Malawi, as part of a study of longitudinal carriage of ESBL-producing Enterobacterales, as previously described^16^. Briefly, we recruited three groups of adults (≥ 16 years): i) 225 adults with sepsis in the emergency department of Queen Elizabeth Central Hospital (QECH), Blantyre, Malawi, ii) 100 antimicrobial-unexposed adults admitted to QECH and iii) 100 antimicrobial-unexposed community dwelling adults. Antimicrobial-unexposed was defined as with no receipt of antimicrobials (except for long-term co-trimoxazole preventative therapy, CPT, or antituberculous chemotherapy) in the previous four weeks. Up to five stool samples per participant (or rectal swab samples performed by trained study team members if participants were unable to provide stool) were collected over the course of six months and aerobically cultured overnight at 37°C on ChromAGAR ESBL-selective chromogenic media (ChromAGAR, France) before being speciated with the API system (Biomeriuex, France).

DNA was extracted from a subsample of isolates identified as *K. pneumoniae* and sequenced: one *K. pneumoniae* colony pick from the first 217 samples where *K. pneumoniae* was identified. DNA was extracted from overnight nutrient broth cultures using the Qiagen DNA Mini kit (Qiagen, Germany) as per the manufacturer’s instructions. DNA was shipped at ambient to the Wellcome Sanger Institute for paired-end 150bp sequencing on the Illumina HiSeq X10 instrument. Species was confirmed with Kraken v1.1.1 and Bracken v2.5 (with a 8Gb MiniKraken database constructed on 3 April 2018).^17^ *De novo* assembly was undertaken with SPAdes v3.10.0^18^ followed by the pipeline by Page et al^19^ and the quality of the assemblies assessed with QUAST v5.0.2^20^ and CheckM v1.1.2^21^; assembly failures with a total assembled length of < 4Mb or assemblies with a CheckM-defined contamination of ≥ 10% were excluded from further analysis. 203 genomes passed QC and were analysed further. Assemblies were then annotated with Prokka v1.14.5 with a genus-specific database from RefSeq^22^ and the Roary v3.13 pangenome pipeline^23^ used to identify core genes with default settings and paralogs not split. Genes present in ≥ 99% samples were considered to be core. 20,853 genes were identified, of which 3391 were core. These were concatenated to a 2.82Mb pseudosequence; the 378,596 variable sites were extracted with SNP-sites v2.5.1^24^ and used to infer the a maximum-likelihood phylogeny with IQ-TREE v1.6.3^25^, with 1000 ultrafast bootstrap replicates. The IQ-TREE ModelFinder module was used to select the best fitting nucleotide substitution model, using ascertainment bias correction; this selected a general time reversible model with FreeRate site heterogeneity and 8 parameters. Trees were visualized using the R *ggtree* v2.2.4^26^ package.

ARIBA v.2.14.6^27^ was used to identify AMR-associated genes using the SRST2 curated version of the ARG-ANNOT database^28^ and to call SNPs in the quinolone-resistance determining regions (QRDR) *gyrA, gyrB, parC* and *parE*, using the wild-type genes from the *Escherichia coli* K-12 substr. MG1655 (NC_000913.3) as reference. Quinolone resistance was assumed to be conferred by QRDR mutations recorded in the Comprehensive Antibiotic Resistance Database^29^ (CARD) as causing quinolone resistance in Enterobacterales. Beta-lactamases were considered to be extended spectrum based on the phenotypic classifications at https://ftp.ncbi.nlm.nih.gov/pathogen/betalactamases/Allele.tab. We explored clustering of AMR genes using hierarchical clustered heatmaps of Jaccard distances of AMR gene presence using the base *dist* and *hclust* functions in R, visualized with the pheatmap package v1.0.2. ARIBA was also used to determine multilocus sequence type (ST) as defined by the 7-gene scheme^30^ hosted at pubMLST (https://pubmlst.org/). Kleborate v2.0.1^31^ was used to infer *Klebsiella* species, capsule polysaccharide (K-type) and lipopolysaccharide (O-type) serotypes, and to identify the presence of the siderophore virulence loci *ybt* (yersiniabactin), *iuc* (aerobactin) and *iro* (salmochelin), the genotoxin locus *clb* (colibactin), and the hypermucoidy genes *rmpA* and *rmpA2*. K- and O-types were recoded as “unknown” if the Kleborate-defined confidence in their identification was below “good”.

To place the isolates from this study in a local and global phylogenetic context, two further phylogenies were constructed, using collections of Malawian isolates^32,33^, from a multi-country, large-scale description of *Klebsiella* population structure^5^ that was designed to span the diversity of the species with samples collected between 1973 and 2011, and 66 genomes from a study of the genomic epidemiology of *Klebsiella* in Kenya^34^. Two studies from QECH provided the previously published Malawian genomes: a genomic investigation into a *K. pneumoniae* outbreak on the neonatal ward at QECH^32^ which sequenced 100 bloodstream infection isolates from children between 2012 and 2015, and a study which sequenced 72 sterile site (blood and CSF) and rectal carriage isolates selected to maximise diversity^33^ from QECH from 1996-2014. The same quality control steps, assembly, annotation and pangenome determination steps were applied; following QC, 150 genomes from samples collected in Malawi were combined with the 203 from this study. Roary identified a pangenome in this Malawian collection of 20,853 of which 3391 were core. These core genes formed a 2.82Mb alignment with 378,596 variable sites which were used to infer a phylogeny for Malawian isolates as described above, using the same nucleotide substitution model. To build a global phylogeny we included all Malawian isolates plus 66 genomes from Kenya^34^ plus 288 genomes from the multi-country^5^ collection, again using the same methods. In total, following QC, this analysis included 687 genomes; the pan-genome as constructed with Roary comprised 49,385 genes, of which 2754 were core; these formed a concatenated pseudosequence of 0.95Mb with 200,622 variable sites, which were used to infer a phylogeny as above; ST and ESBL presence or absence were also inferred as described above.

All statistical analyses were carried out in R v4.0.2 (R Foundation for Statistical Computing, Vienna, Austria). Summary statistics, where presented, are medians and interquartile ranges or proportions with exact binomial confidence intervals unless otherwise stated. Comparisons of proportions use Fisher’s exact test; to compare proportions of each O- and K-type between invasive and colonising isolates we corrected for multiple comparisons using the Benjamini-Hochberg p-value correction. The clinical study which provided the isolates for this analysis was approved by the Liverpool School of Tropical Medicine (16-062) and Malawi College of Medicine (P.11/16/2063) research ethics committees. The code and data to reproduce the analyses in this manuscript are available as the *blantyreESBL* v1.0.0^35^ R package on GitHub at https://joelewis101.github.io/blantyreESBL/. Reads from all isolates sequenced as part of this study have been deposited in the European Nucleotide Archive, and accession numbers (as well as accession numbers of publicly available genomes used in this analysis) are provided in the R package.

## Results

### Population structure

Most of the 203 included *K. pneumoniae* complex genomes sequenced for this study were *Klebsiella pneumoniae subsp. pneumoniae* (n=190), *Klebsiella quasipneumoniae subsp. similipneumoniae* (n=7*), Klebsiella variicola subsp. variicola* (n=3), *Klebsiella quasipneumoniae subsp. quasipneumoniae* (n=2) and *Klebsiella quasivariicola* (n=1) were also represented, comparable to other clinical studies on the *K. pneumoniae* species complex^36^. We identified a large number of STs considering the number of samples (Figure 1) with 61 identified and a median of 2 (IQR 1-4) isolates per ST; 26/61 STs were only represented once in the collection. The most common STs identified were ST307 (n=16), ST14 (n= 14), ST340 (n=12), and ST15 (n=11). The STs reflected the structure of the inferred core-gene phylogeny well (Supplementary Figure 1), which, as expected, had the characteristic *Klebsiella* deep branching topology^2^.

**Figure 1:**
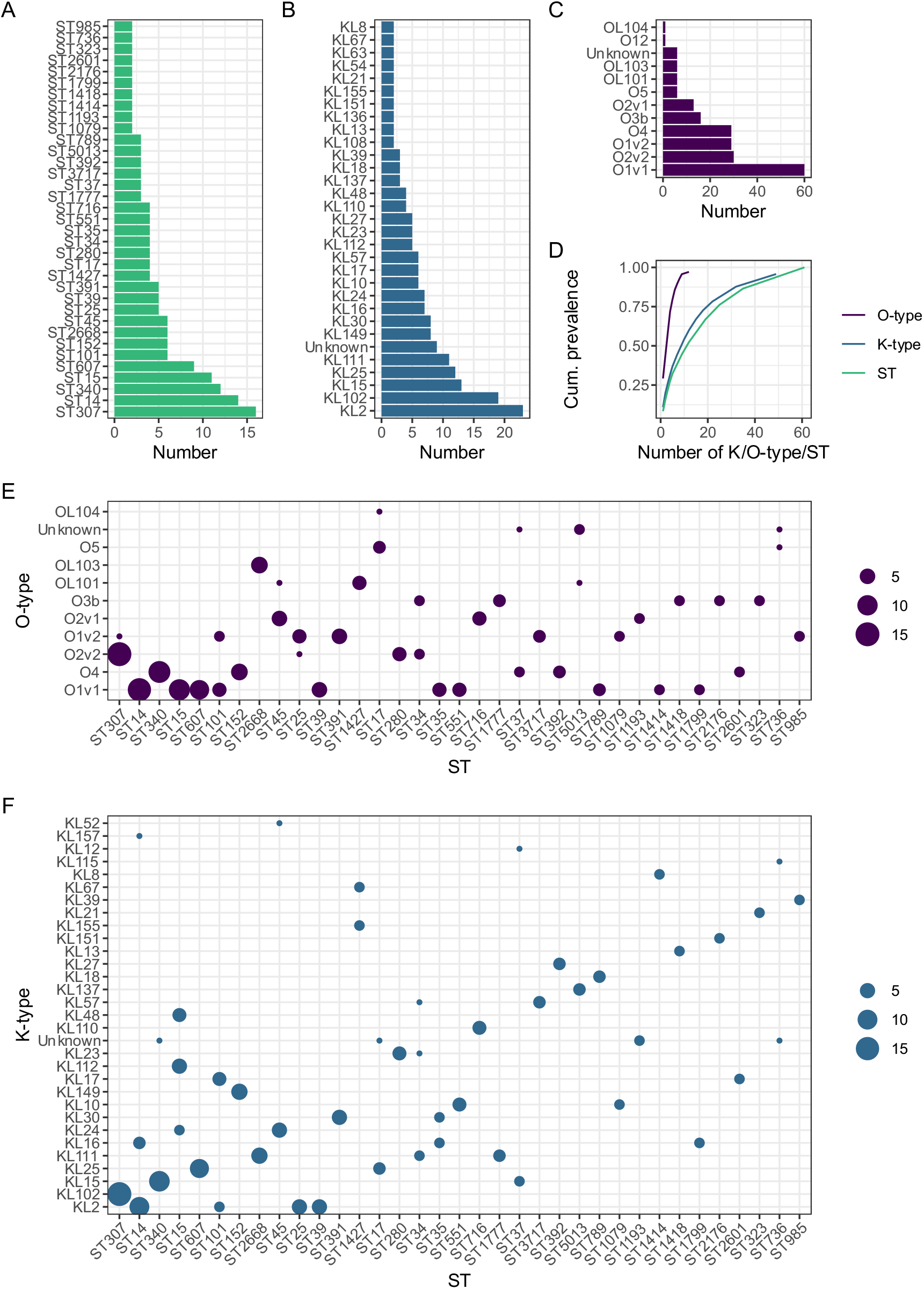
Diversity of *K. pneumoniae* included in the study. Distributions of (A) sequence type (ST) (B) K-type (C) O-type. (D) shows cumulative prevalence as a function of number of K/O-types or ST, where K/O-type or ST is ordered from largest to smallest. (E) and (F) show ST association of O-type and K-type respectively, where area of point is proportional to number of samples. STs with only a single representative in the collection are excluded from plots A, B, C, E, F.

Adding Malawian invasive isolates (n=150) to the Malawian colonising isolates (n=203) showed that they had a similar population structure (Figure 2). The global core-gene based phylogeny including global context genomes and Malawian invasive and colonizing isolates showed that Malawian isolates were distributed throughout the tree (Figure 3); Malawian human colonising and invasive isolates mirror the broader *Klebsiella* population structure of the included samples. The ST14 and ST15 isolates clustered with genomes from elsewhere but ST340 and ST307 were Malawi-restricted.

**Figure 2:**
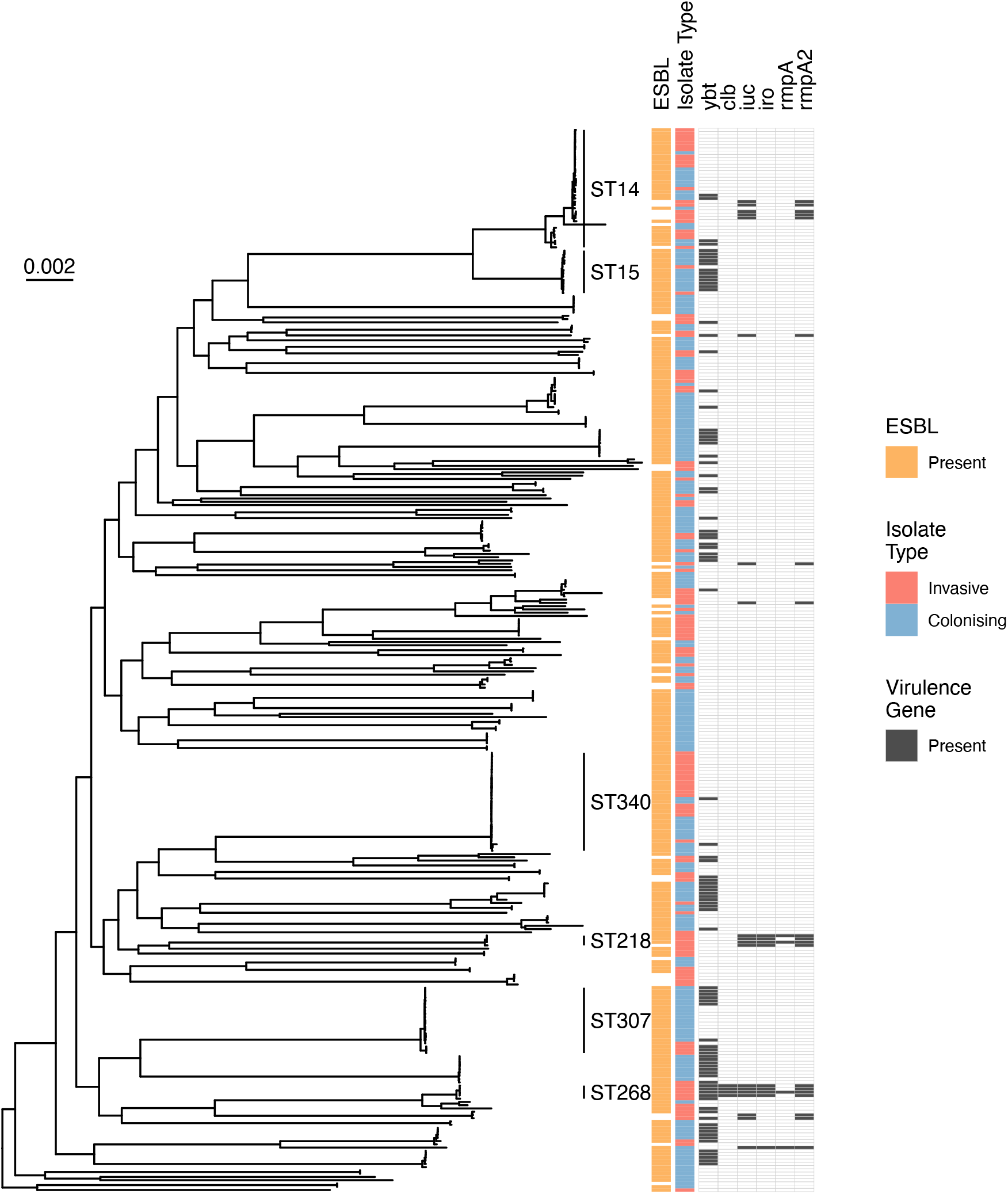
Midpoint rooted core-gene maximum-likelihood phylogenetic tree of Malawian isolates, including all genomes from this study and context genomes from Malawian studies (n = 353), and restricted to *Klebsiella pneumoniae sensu stricto* (KpI). Heatmaps show whether ESBL genes are present, whether carriage or infection, and whether the siderophore virulence loci *ybt* (yersinobactin), *iuc* (aerobactin), *iro* (salmochein), the genotoxin virulence locus *clb* (colibactin) and the hypermucoidy genes *rmpA* and *rmpA2* are present. Scale bar shows nucleotide substitutions per site.

**Figure 3:**
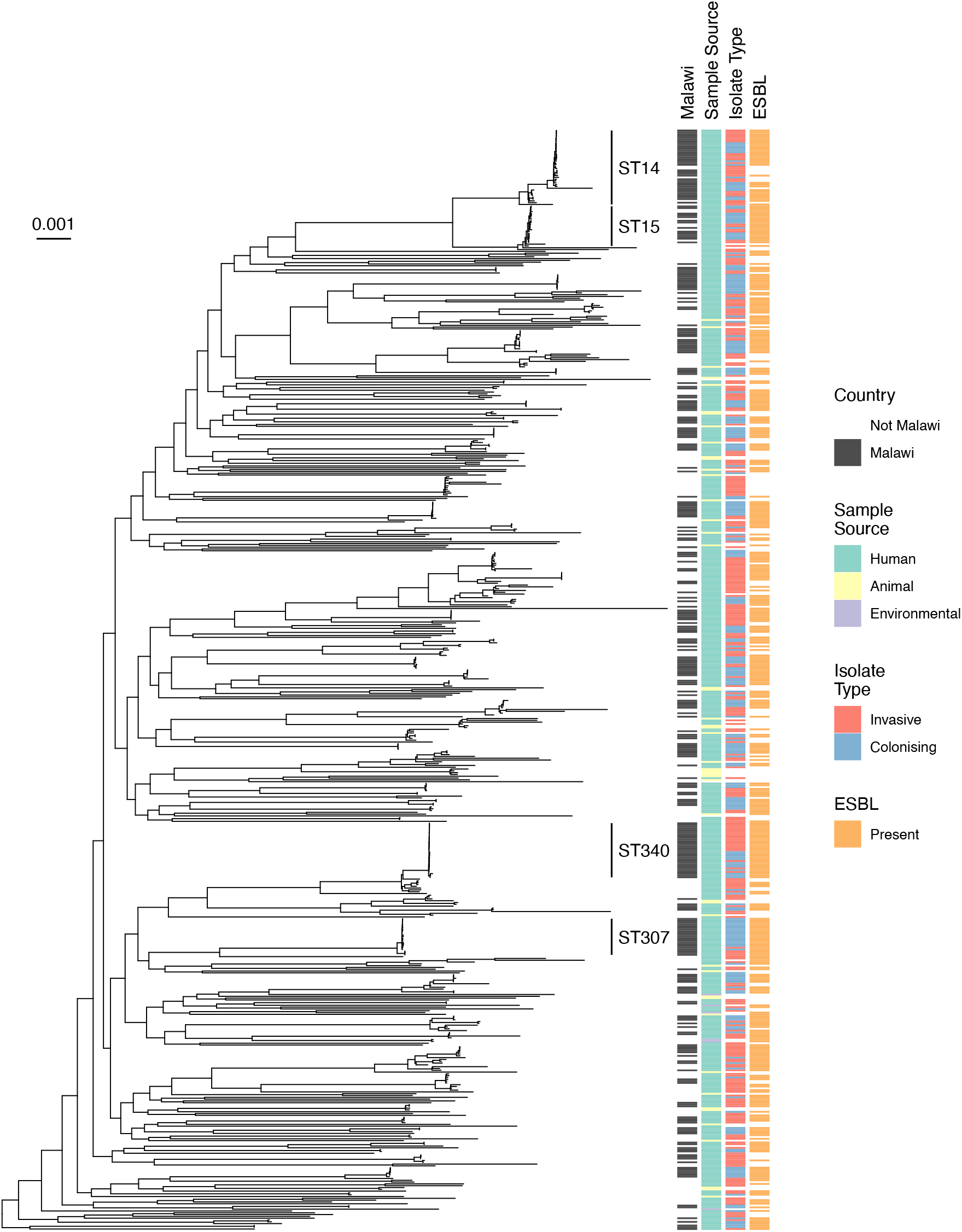
Midpoint rooted core-gene phylogenetic tree of Malawian and global isolates, restricted to *Klebsiella pneumoniae* sensu stricto (KpI). Heatmaps show whether isolated in Malawi, whether animal/human/environmental, whether carriage or infection, and whether ESBL genes are present. Scale bar shows nucleotide substitutions per site.

### Antigenic and AMR gene diversity of Malawian colonising isolates

49 K-types were identified and were ST-associated with median 1 (IQR 1-2) STs per K-type. Eight O-types were identified: six of the 12 originally described O-types based on serology (O1 [v1 and v2], O2 [v1 and v2], 3b, 4, 5, 12), as well as 3 *rfb* loci predicted to encode novel O-types which have been reported but not yet described serologically^37^, denoted by OL: OL101, 103, and 104, though these were in the minority (13/203; Figure 1). O-types were more likely to be encoded by multiple STs than K types: each O-type was associated with a median of six (IQR 2.5 – 11.8) STs. The four most common predicted O-antigens were O1 (89/203 [44%]), O2 (43/203 [21%]), O4 (29/203 [14%]) and O3b (16/203 [8%]) accounting for 87% of samples (Supplementary Table 1). In contrast, the four most frequent K-types (K2 [11%], 102 [9%], 15 [6%] and 25 [6%], Supplementary Table 2) and STs (ST307 [8%], 14 [7%], 340 [6%], 15 [5%]) together accounted for 33% and 26% of samples, respectively.

The isolates contained a median of 15 (IQR 12-17, range 6-25) antimicrobial resistance genes (including SNPs) per genome (Figure 4). Consistent with isolates being grown on ESBL selective media, at least one ESBL-encoding gene was identified in 99% of genomes (200/203); genes encoding narrow spectrum beta-lactamases were also common (200/203, 99% genomes, excluding the genus-associated *bla*_ampH_ penicillinase which was present in 100% of isolates). The most commonly identified ESBL-encoding gene was *bla*_CTX-M-15_ in 186/203 (92%) of genomes. Genes conferring resistance to sulphonamides (201/203, 99%), trimethoprim (198/203, 98%) and aminoglycosides (198/203 98%) were near-ubiquitous; determinants of resistance to chloramphenicol (140/203, 69%) and fluoroquinolones (76/203, 37%) were less common. Quinolone resistance determinant region mutations identified were and S83F (n=10) in *gyrA* and S80I (n=1) in *parC*, but plasmid-mediated quinolone resistance determinants *qnrB* (n= 30) and *qnrS* (n=40) were also identified. No known genes conferring carbapenemase resistance were identified.

**Figure 4:**
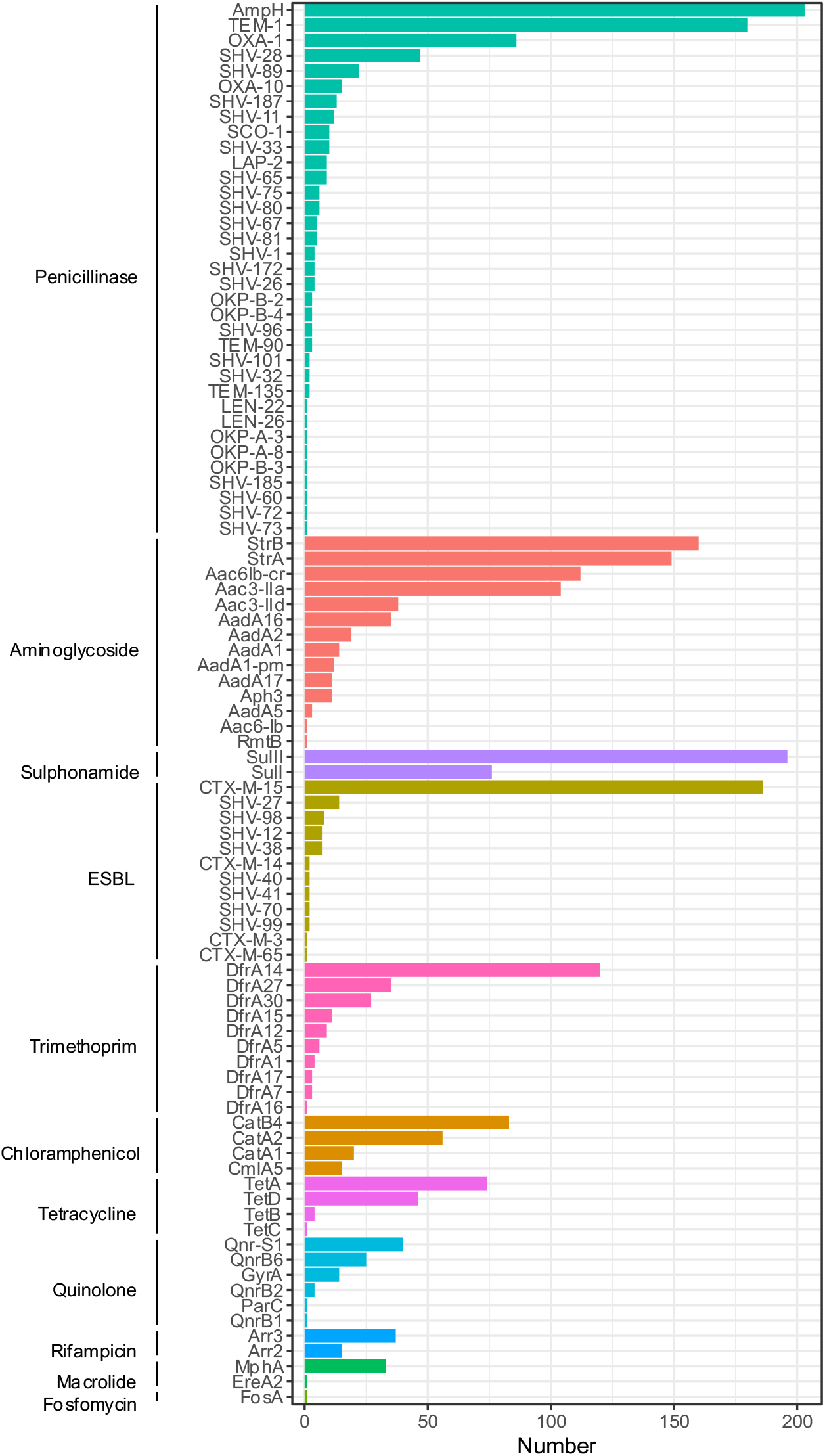
Antimicrobial resistance determinants identified.

Some AMR genes clustered together (e.g. *strA* with *strB, bla*_CTX-M-15_ with *bla*_TEM-1_ and *sulII*, and *bla*_SHV-11_ with *aadA1-pm* and *bla*_OXA-10_, Supplementary Figure 2), and some of these AMR-gene clusters were lineage-associated (e.g. the *bla*_SHV-11_ *aadA1-pm bla*_OXA-10_ cluster with ST340, Supplementary Figure 3).

### Comparing Malawian carriage and bloodstream isolates

Finally, we compared Malawian carriage to invasive isolates in terms of K- and O-type and recognized virulence determinants. O-type and K-type distributions were similar across invasive and colonising isolates (Figure 5). Fisher’s exact tests corrected for multiple comparisons were consistent with similar proportions of each O-type and K-type across infecting and colonizing isolates, with the exception of KL62 and KL43 (Supplementary Tables 3 and 4). Both of these K-types were strongly associated with invasive isolates (9/10 KL62 and 9/9 KL43 were invasive) and associated with multiple STs: KL43 was present in ST372 (6 isolates), ST106 (2 isolates), ST276 (1 isolate) and KL62 was present in ST644 (4 isolates), ST48 (3 isolates), ST348 (2 isolates) and ST4 and ST432 (one isolate each). All of these isolates contained O-type 1 or 2.

**Figure 5:**
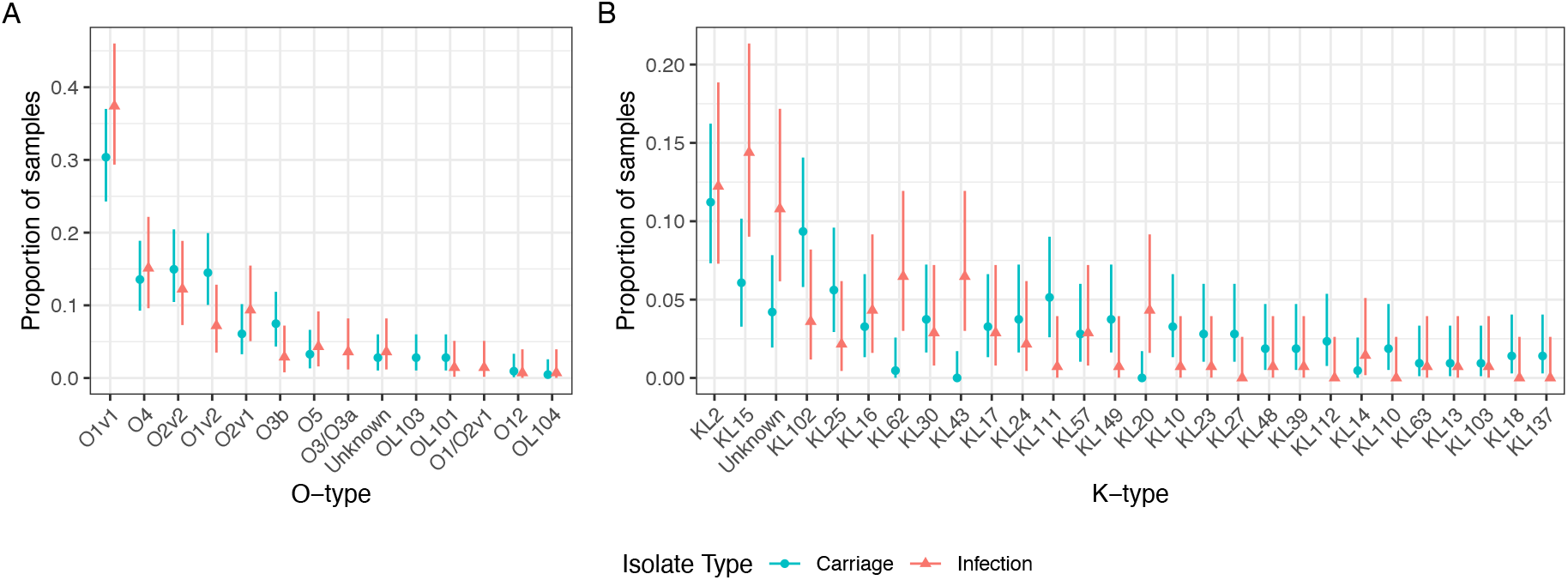
Distribution of O-types (A) and K-types (B) stratified by carriage or infecting samples, showing that the O-type distribution is similar whether infecting of colonizing.

The most commonly identified virulence determinant was the siderophore locus *ybt*, present in 27% (93/365) of Malawian isolates, and more common in carriage (68/214, 32%) compared to invasive (26/139, 19%, p = 0.006) isolates, and in ESBL isolates (89/306, 29%) compared to non-ESBL isolates (5/47, 11%, p = 0.007). All other virulence determinants were less common (the siderophore loci *iuc* in 5% and *iro* in 3%; the genotoxin loci *clb* in 3%; and the hypermucoidy genes *rmpA* and *rmpA2* in 5% and 3% respectively), but were strongly associated with invasive isolates: only 1/214 (0.5%) carriage isolate contained any of these virulence genes (Figure 2 and Supplementary Figure 4), compared to 18/139 of invasive isolates (12.9%, p < 0.001). Generally, isolates containing these non-*ybt* virulence determinants were less likely to contain ESBL-encoding genes (11/47 [23%] non-ESBL isolates vs 8/306 [3%], p < 0.001) but two lineages contained both: ST268 (4 isolates with *ybt, clb, iro,iuc* and *rmpA2* +/- *rmpA*) and ST218 (4 isolates with *iuc, iro* and *rmpA2* +/- *rmpA*) also all carried ESBL-encoding genes. These ST218/268 isolates were all from blood culture, the earliest in 2004 and no isolates of these STs were cultured from carriage samples. All ST218 were identified as K-locus KL57 and ST268 as KL20, both of which have been described as virulence-associated K-types^8^. Of the two K-types which seemed to be associated with invasive diseases in Malawi (KL43 and 62), none of the nine KL43-containing isolates contained any other identified virulence determinant. Of the ten KL62-containing isolates, 4/10 contained no other identified virulence determinant; 4/10 contained *ybt* alone and one contained *ybt* and *iuc*, and one *iuc* and *rmpA*.

## Discussion

Malawi, and sSA in general, is an area of the world that is undersampled in current *K. pneumoniae* collections. We present a genomic investigation of 203 colonising ESBL-producing *K. pneumoniae* species complex genomes from Blantyre. Placing these in both a Malawian and global context enables us to draw several conclusions that increase our understanding of the dynamics of colonisation and infection in this setting, as well as understand the differential threats for the global emergence of drug resistant Klebsiella from all parts of the world.

The isolates causing infection in Malawi and the isolates from carriage represent highly similar population structures, consistent with the hypothesis that the source of infecting *K. pneumoniae* is the host microbiota^1^. Importantly, the diversity seen in the Malawian ESBL *K. pneumoniae* reflects the diversity of *K. pneumoniae* isolates from multiple countries, suggesting that Malawi samples global *K. pneumoniae* diversity. However, there are important local differences: some STs are overrepresented in the Malawian isolates as compared to the multi-country collections we used for context, suggesting different requirements for success (and subsequent clonal expansion) in Malawi as to the other (largely high-income country) settings included. These successful Malawian STs included ST14 and ST15, which are common ESBL and carbapenemase-associated lineages particularly in Asia^8^; Malawian isolates clustered closely with context ST14/15 genomes in the core gene tree. In comparison, the core gene tree topology of the common Malawian ST307 and ST340 lineages was consistent with local subclades. This may be representative of the biases inherent in the sampling frame of the multicounty collection; ST307 has been found to be a common ESBL- and carbapenemase associated lineage in North^14^ and South^38^ America, and is represented in European collections^15^, as is ST340. However, these STs are unusual in Asia^8^, from where many of the multicountry context isolates arise.

The drivers of local or regional success for some lineages over others are not clear; in the Malawian setting, success was not explained by the identified virulence determinants. The well described virulence loci *clb, ioc, iuc* and hypermucoidy genes *rmpA* ad *rmpA2* are, as expected, present in multiple STs causing invasive disease in the Malawi. Although presence of these loci was associated with invasive disease, the majority of invasive isolates lacked them, suggesting that host factors or other pathogen factors (e.g. capsule) are the primary determinant of whether carriage develops into infection in the Malawian setting. Of note, the K-types KL62 and KL43 were associated with invasive disease in our setting. Generally, presence of virulence determinants was associated with absence of ESBL-encoding genes, as described elsewhere in the world^2^; however, two lineages (ST218 and ST268) demonstrated the presence of both, indicating that hypervirulent-AMR Klebsiella lineages are present in Malawi.

AMR-gene content was diverse with median 15 AMR genes per isolate, consistent with previous studies^36^; *bla*_CTX-M-15_ was by far the most commonly identified ESBL-encoding gene, present in 92% of isolates. Presence of trimethoprim and sulphonamide resistance determinants was near universal. In Malawi, a high-HIV prevalence setting, the WHO recommends lifelong co-trimoxazole preventative therapy for people living with HIV,^39^ and it is also consequently widely used as a mainstay of community antimicrobial chemotherapy in health centres in Malawi (source: Ministry of Health Malawi). This raises the possibility that co-location of genes conferring ESBL and co-trimoxazole could select for ESBL resistance even in the absence of beta-lactam use. No genes conferring resistance to carbapenemases were identified, though the carbapenemase *bla*_NDM-5_ has been described in *E. coli* in Blantyre contemporaneously with this study^40^, and the carbapenemases *bla*_KPC-2 in_ ST340 *K. pneumoniae* and *bla*_OXA-48_ in *K. variicola* have been described in Central Malawi^41^ in 2016/17. Carbapenems are increasingly available in QECH, and it seems very likely that increasing exposure will result in rapid expansion of carbapenemases in *K. pneumoniae* especially given the likely unrestricted global flow of Klebsiella strains suggested by our data.

There was significant diversity in the Malawian collection in K- and O-types, as is characteristic of Klebsiella collections^8,13,42^. Vaccines based on *K. pneumoniae* O antigens are in development^10^, and quadrivalent vaccines with various O-antigen targets have been proposed, depending on the cohort used to describe O-antigen epidemiology: O1, 2, 3 and 5 would have activity against 90% of *K. pneumoniae* strains, in one global collection of 645 isolates;^13^ a second study suggested O2v2, O1v1, O3b, O1v2 would cover 71-77% of European isolates, based on a derivation isolate collection from Oxfordshire, UK^42^. O-antigen variation is apparent depending on location of isolation: the global collection described above contained 11% O5 containing isolates, but the Oxfordshire collection 3%, as did our collection. In the Malawian setting, O4 was identified in 14% of carriage and 15% of infecting isolates, but was rare in the global (2%)^13^ and Oxfordshire (4%)^42^ collections and also in South East Asia^8^ (2-4%). This highlights the need for longitudinal surveillance and truly global collections, describing secular trends in O- and K-antigen epidemiology from diverse settings to guide vaccine development. We found that the O-type distribution for Malawian ESBL producing *K. pneumoniae* carriage isolates was similar to invasive isolates, suggesting that stool or rectal swab sampling with selective culture could be a cost-effective way to rapidly expand understanding of worldwide O-type distributions to guide vaccine development. This finding must be confirmed in further sites before such a strategy could be adopted.

There are limitations to our study. Most importantly, our sampling scheme is not unbiased. ESBL-producing carriage isolates were selected for, and one of the Malawian studies providing invasive context genomes was an investigation of a *K. pneumoniae* outbreak on the QECH neonatal unit. This is likely to have introduced bias into the collection of Malawian genomes, especially against classically hypervirulent but antimicrobial susceptible lineages. All Malawian genomes are from a single centre, which enables us to compare the population structure of carriage and clinical isolates but may limit generalisation to other settings in sSA. Multiple samples were cultured from single individuals and so were not independent, which could introduce bias, however most individuals were colonized by different strains at different time points

In conclusion, we present a genomic analysis of ESBL *K. pneumoniae* colonising adults in Blantyre, Malawi. Malawian colonizing and invasive isolates are similar and population structure is comparable to global *Klebsiella* population structure. This suggests that Malawi is sampling global Klebsiella diversity - but we demonstrate that some lineages (ST14, ST15, ST307, ST340) are successful in the Malawian setting and have undergone expansion in this setting. The reason for this success is not explained by the virulence factors we sought and host factors, pathogen opportunity, or alternate virulence factors not linked to disease elsewhere may be the major determinants of lineage success and invasive disease in Malawi. O-antigen distributions of Malawian isolates showed some differences to previously described collections, highlighting the need for geographically-aware surveillance to inform vaccine development. Predicted O-antigen diversity was similar across invasive and carriage isolates, suggesting that O-typing of carriage isolates could be a costeffective way to rapidly carry out such surveillance and assess putative O-antigen vaccine coverage across diverse populations.

## Supporting information

Supplementary Results

## Author contributions

Conceptualisation: JL, NT, NAF. Methodology: JL, NT, NAF, MAB, EH, JM. Investigation: JL, MM, RB. Formal analysis: JL, NT, NAF, EH, MAB. Writing – original draft preparation; JL. Writing – review and editing: JL,MM,RB, MB,JM,EH,NT,NAF. Supervision: NAF,NT

## Acknowledgements

Many thanks to the study team: Lucy Keyala, Tusekile Phiri, Grace Mwaminawa, Witness Mtambo, Gladys Namacha, Monica Matola; to the MLW laboratory teams, particularly Brigitte Denis; and to the MLW data team, particularly Lumbani Makhaza and Clemens Masesa. The authors acknowledge the sequencing team at the Wellcome Sanger Institute, and Christoph Puethe and the Pathogen Informatics team for computational support.

## References

1. Gorrie, C. L. et al. Gastrointestinal Carriage Is a Major Reservoir of Klebsiella pneumoniae Infection in Intensive Care Patients. Clinical Infectious Diseases 65, 208–215 (2017).

2. Wyres, K. L., Lam, M. M. C. & Holt, K. E. Population genomics of Klebsiella pneumoniae. Nat Rev Microbiol 18, 344–359 (2020).

3. World Health Organisation. Prioritization of pathogens to guide discovery, research and development of new antibiotics for drug-resistant bacterial infections, including tuberculosis. (2017).

4. Musicha, P. et al. Trends in antimicrobial resistance in bloodstream infection isolates at a large urban hospital in Malawi (1998–2016): a surveillance study. The Lancet Infectious Diseases 17, 1042–1052 (2017).

5. Holt, K. E. et al. Genomic analysis of diversity, population structure, virulence, and antimicrobial resistance in Klebsiella pneumoniae, an urgent threat to public health. PNAS 112, E3574–E3581 (2015).

6. Lam, M. M. C. et al. Genetic diversity, mobilisation and spread of the yersiniabactin-encoding mobile element ICEKp in Klebsiella pneumoniae populations. Microbial Genomics, 4, e000196 (2018).

7. Walker, K. A. et al. A Klebsiella pneumoniae Regulatory Mutant Has Reduced Capsule Expression but Retains Hypermucoviscosity. mBio 10, (2019).

8. Wyres, K. L. et al. Genomic surveillance for hypervirulence and multi-drug resistance in invasive Klebsiella pneumoniae from South and Southeast Asia. Genome Medicine 12, 11 (2020).

9. Russo, T. A. & Marr, C. M. Hypervirulent Klebsiella pneumoniae. Clinical Microbiology Reviews 32, e00001–19 (2019).

10. Choi, M., Tennant, S. M., Simon, R. & Cross, A. S. Progress towards the development of Klebsiella vaccines. Expert Review of Vaccines 18, 681–691 (2019).

11. Follador, R. et al. The diversity of Klebsiella pneumoniae surface polysaccharides. Microbial Genomics, 2, e000073 (2016).

12. Wyres, K. L. et al. Identification of Klebsiella capsule synthesis loci from whole genome data. Microbial Genomics 2, e000102.

13. Choi, M. et al. The Diversity of Lipopolysaccharide (O) and Capsular Polysaccharide (K) Antigens of Invasive Klebsiella pneumoniae in a Multi-Country Collection. Front Microbiol 11, (2020).

14. Long, S. W. et al. Population Genomic Analysis of 1,777 Extended-Spectrum Beta-Lactamase-Producing Klebsiella pneumoniae Isolates, Houston, Texas: Unexpected Abundance of Clonal Group 307. mBio 8, e00489–17 (2017).

15. David, S. et al. Epidemic of carbapenem-resistant Klebsiella pneumoniae in Europe is driven by nosocomial spread. Nat Microbiol 4, 1919–1929 (2019).

16. Lewis, J. et al. Dynamics of gut mucosal colonisation with extended spectrum beta-lactamase producing Enterobacterales in Malawi [submitted]. bioRxiv.

17. Wood, D. E. & Salzberg, S. L. Kraken: ultrafast metagenomic sequence classification using exact alignments. Genome Biology 15, R46 (2014).

18. Bankevich, A. et al. SPAdes: a new genome assembly algorithm and its applications to single-cell sequencing. Journal of computational biology : a journal of computational molecular cell biology 19, 455–77 (2012).

19. Page, A. J. et al. Robust high-throughput prokaryote de novo assembly and improvement pipeline for Illumina data. Microbial Genomics 2, e000083.

20. Gurevich, A., Saveliev, V., Vyahhi, N. & Tesler, G. QUAST: quality assessment tool for genome assemblies. Bioinformatics 29, 1072–1075 (2013).

21. Parks, D. H., Imelfort, M., Skennerton, C. T., Hugenholtz, P. & Tyson, G. W. CheckM: assessing the quality of microbial genomes recovered from isolates, single cells, and metagenomes. Genome research 25, 1043–55 (2015).

22. Seemann, T. Prokka: rapid prokaryotic genome annotation. Bioinformatics 30, 2068–2069 (2014).

23. Page, A. J. et al. Roary: rapid large-scale prokaryote pan genome analysis. Bioinformatics 31, 3691–3693 (2015).

24. Page, A. J. et al. SNP-sites: rapid efficient extraction of SNPs from multi-FASTA alignments. Microbial Genomics 2, e000056 (2016).

25. Nguyen, L.-T., Schmidt, H. A., von Haeseler, A. & Minh, B. Q. IQ-TREE: A Fast and Effective Stochastic Algorithm for Estimating Maximum-Likelihood Phylogenies. Molecular Biology and Evolution 32, 268–274 (2015).

26. Yu, G., Smith, D. K., Zhu, H., Guan, Y. & Lam, T. T.-Y. ggtree : an r package for visualization and annotation of phylogenetic trees with their covariates and other associated data. Methods in Ecology and Evolution 8, 28–36 (2017).

27. Hunt, M. et al. ARIBA: rapid antimicrobial resistance genotyping directly from sequencing reads. Microbial genomics 3, e000131 (2017).

28. Inouye, M. et al. SRST2: Rapid genomic surveillance for public health and hospital microbiology labs. Genome Medicine 6, 90 (2014).

29. Alcock, B. P. et al. CARD 2020: antibiotic resistome surveillance with the comprehensive antibiotic resistance database. Nucleic Acids Res 48, D517–D525 (2020).

30. Diancourt, L., Passet, V., Verhoef, J., Grimont, P. A. D. & Brisse, S. Multilocus Sequence Typing of Klebsiella pneumoniae Nosocomial Isolates. Journal of Clinical Microbiology 43, 4178–4182 (2005).

31. Lam, M. M. C. et al. A genomic surveillance framework and genotyping tool for Klebsiella pneumoniae and its related species complex. Nat Commun 12, 4188 (2021).

32. Cornick, J. et al. Genomic investigation of a suspected multi-drug resistant Klebsiella pneumoniae outbreak in a neonatal care unit in sub-Saharan Africa. bioRxiv 2020.08.06.236117 (2020) doi:10.1101/2020.08.06.236117.

33. Musicha, P. et al. Genomic analysis of Klebsiella pneumoniae isolates from Malawi reveals acquisition of multiple ESBL determinants across diverse lineages. Journal of Antimicrobial Chemotherapy 74, 1223–1232 (2019).

34. Henson, S. P. et al. Molecular epidemiology of Klebsiella pneumoniae invasive infections over a decade at Kilifi County Hospital in Kenya. Int J Med Microbiol 307, 422–429 (2017).

35. Lewis, J. joelewis101/blantyreESBL: v1.0.0. (Zenodo, 2021). doi:10.5281/zenodo.5554082.

36. Ellington, M. J. et al. Contrasting patterns of longitudinal population dynamics and antimicrobial resistance mechanisms in two priority bacterial pathogens over 7 years in a single center. Genome Biology 20, 184 (2019).

37. Kaptive Web: User-Friendly Capsule and Lipopolysaccharide Serotype Prediction for Klebsiella Genomes | Journal of Clinical Microbiology. https://jcm.asm.org/content/56/6/e00197-18.

38. Ocampo, A. M. et al. A Two-Year Surveillance in Five Colombian Tertiary Care Hospitals Reveals High Frequency of Non-CG258 Clones of Carbapenem-Resistant Klebsiella pneumoniae with Distinct Clinical Characteristics. Antimicrobial Agents and Chemotherapy 60, 332–342.

39. World Health Organisation. Consolidated guidelines on the use of antiretroviral drugs for treating and preventing HIV infection: recommendations for a public health approach. Second edition. (2016).

40. Lewis, J. M. et al. Emergence of carbapenemase producing Enterobacteriaceae, Malawi. J Glob Antimicrob Resist 20, 225–227 (2019).

41. Kumwenda, G. P. et al. First Identification and genomic characterization of multidrug-resistant carbapenemase-producing Enterobacteriaceae clinical isolates in Malawi, Africa. J Med Microbiol 68, 1707–1715 (2019).

42. Lipworth, S. et al. Ten years of population-level genomic Escherichia coli and Klebsiella pneumoniae serotype surveillance informs vaccine development for invasive infections. Clinical Infectious Diseases (2021) doi:10.1093/cid/ciab006.

