## Supplementary Results for "Genomic and antigenic diversity of carried *Klebsiella pneumoniae* isolates mirrors that of invasive isolates in Blantyre, Malawi"

### Supplementary Material

**Supplementary Table 1:** Distribution of O-antigens

| O-type | n (%) | Cumulative n (%) |
|---|---|---|
| O1v1 | 60 (29.6%) | 60 (29.6%) |
| O2v2 | 30 (14.8%) | 90 (44.3%) |
| O1v2 | 29 (14.3%) | 119 (58.6%) |
| O4 | 29 (14.3%) | 148 (72.9%) |
| O3b | 16 (7.9%) | 164 (80.8%) |
| O2v1 | 13 (6.4%) | 177 (87.2%) |
| O5 | 6 (3.0%) | 183 (90.1%) |
| OL101 | 6 (3.0%) | 189 (93.1%) |
| OL103 | 6 (3.0%) | 195 (96.1%) |
| Unknown | 6 (3.0%) | 201 (99.0%) |
| O12 | 1 (0.5%) | 202 (99.5%) |
| OL104 | 1 (0.5%) | 203 (100.0%) |

**Supplementary Table 2:** Distribution of K-antigens

| K-type | n (%) | Cumulative n (%) |
|---|---|---|
| KL2 | 23 (11.3%) | 23 (11.3%) |
| KL102 | 19 (9.4%) | 42 (20.7%) |
| KL15 | 13 (6.4%) | 55 (27.1%) |
| KL25 | 12 (5.9%) | 67 (33.0%) |
| KL111 | 11 (5.4%) | 78 (38.4%) |
| Unknown | 9 (4.4%) | 87 (42.9%) |
| KL149 | 8 (3.9%) | 95 (46.8%) |
| KL30 | 8 (3.9%) | 103 (50.7%) |
| KL16 | 7 (3.4%) | 110 (54.2%) |
| KL24 | 7 (3.4%) | 117 (57.6%) |
| KL10 | 6 (3.0%) | 123 (60.6%) |
| KL17 | 6 (3.0%) | 129 (63.5%) |
| KL57 | 6 (3.0%) | 135 (66.5%) |
| KL112 | 5 (2.5%) | 140 (69.0%) |
| KL23 | 5 (2.5%) | 145 (71.4%) |
| KL27 | 5 (2.5%) | 150 (73.9%) |
| KL110 | 4 (2.0%) | 154 (75.9%) |
| KL48 | 4 (2.0%) | 158 (77.8%) |
| KL137 | 3 (1.5%) | 161 (79.3%) |
| KL18 | 3 (1.5%) | 164 (80.8%) |
| KL39 | 3 (1.5%) | 167 (82.3%) |
| KL108 | 2 (1.0%) | 169 (83.3%) |
| KL13 | 2 (1.0%) | 171 (84.2%) |
| KL136 | 2 (1.0%) | 173 (85.2%) |
| KL151 | 2 (1.0%) | 175 (86.2%) |
| KL155 | 2 (1.0%) | 177 (87.2%) |
| KL21 | 2 (1.0%) | 179 (88.2%) |
| KL54 | 2 (1.0%) | 181 (89.2%) |
| KL63 | 2 (1.0%) | 183 (90.1%) |
| KL67 | 2 (1.0%) | 185 (91.1%) |
| KL8 | 2 (1.0%) | 187 (92.1%) |
| KL103 | 1 (0.5%) | 188 (92.6%) |
| KL105 | 1 (0.5%) | 189 (93.1%) |
| KL114 | 1 (0.5%) | 190 (93.6%) |
| KL115 | 1 (0.5%) | 191 (94.1%) |
| KL12 | 1 (0.5%) | 192 (94.6%) |
| KL123 | 1 (0.5%) | 193 (95.1%) |
| KL14 | 1 (0.5%) | 194 (95.6%) |
| KL145 | 1 (0.5%) | 195 (96.1%) |
| KL157 | 1 (0.5%) | 196 (96.6%) |
| KL19 | 1 (0.5%) | 197 (97.0%) |
| KL52 | 1 (0.5%) | 198 (97.5%) |
| KL59 | 1 (0.5%) | 199 (98.0%) |
| KL62 | 1 (0.5%) | 200 (98.5%) |
| KL64 | 1 (0.5%) | 201 (99.0%) |
| KL74 | 1 (0.5%) | 202 (99.5%) |
| KL9 | 1 (0.5%) | 203 (100.0%) |

**<u>Supplementary Table 3:</u>** Fisher's exact test Benjamini-Hochberg corrected p-values testing equal distribution of O-types across invasive and carriage isolates

| **O-type** | **Uncorrected p-value** | **Benjamini-Hochberg corrected p-values** |
|---|---|---|
| O1/O2v1 | 0.15000 | 0.420 |
| O12 | 1.00000 | 1.000 |
| O1v1 | 0.20000 | 0.467 |
| O1v2 | 0.05960 | 0.342 |
| O2v1 | 0.29600 | 0.592 |
| O2v2 | 0.63600 | 0.897 |
| O3/O3a | 0.00846 | 0.118 |
| O3b | 0.09780 | 0.342 |
| O4 | 0.64100 | 0.897 |
| O5 | 1.00000 | 1.000 |
| OL101 | 0.49100 | 0.859 |
| OL103 | 0.08550 | 0.342 |
| OL104 | 1.00000 | 1.000 |
| Unknown | 0.75600 | 0.962 |

**Supplementary Table 4:** Fisher's exact test Benjamini-Hochberg corrected p-values testing equal distribution of K-types across invasive and carriage isolates

| K-type | **Uncorrected p-value** | **Benjamini-Hochberg corrected p-values** |
|---|---|---|
| KL10 | 0.157000 | 0.6680 |
| KL102 | 0.054800 | 0.6030 |
| KL103 | 1.000000 | 1.0000 |
| KL104 | 0.389000 | 0.8280 |
| KL105 | 1.000000 | 1.0000 |
| KL106 | 1.000000 | 1.0000 |
| KL108 | 0.523000 | 0.8850 |
| KL109 | 0.389000 | 0.8280 |
| KL110 | 0.160000 | 0.6680 |
| KL111 | 0.032900 | 0.4340 |
| KL112 | 0.161000 | 0.6680 |
| KL114 | 1.000000 | 1.0000 |
| KL115 | 1.000000 | 1.0000 |
| KL12 | 1.000000 | 1.0000 |
| KL122 | 0.389000 | 0.8280 |
| KL123 | 1.000000 | 1.0000 |
| KL125 | 0.389000 | 0.8280 |
| KL127 | 0.389000 | 0.8280 |
| KL13 | 1.000000 | 1.0000 |
| KL132 | 1.000000 | 1.0000 |
| KL134 | 0.389000 | 0.8280 |
| KL136 | 0.523000 | 0.8850 |
| KL137 | 0.285000 | 0.8280 |
| KL14 | 0.562000 | 0.9050 |
| KL142 | 0.389000 | 0.8280 |
| KL145 | 1.000000 | 1.0000 |
| KL149 | 0.162000 | 0.6680 |
| KL15 | 0.008640 | 0.1430 |
| KL151 | 0.523000 | 0.8850 |
| KL155 | 0.523000 | 0.8850 |

| | | |
|---|---|---|
| KL157 | 1.000000 | 1.0000 |
| KL158 | 0.389000 | 0.8280 |
| KL16 | 0.576000 | 0.9050 |
| KL165 | 0.389000 | 0.8280 |
| KL17 | 1.000000 | 1.0000 |
| KL18 | 0.285000 | 0.8280 |
| KL19 | 1.000000 | 1.0000 |
| KL2 | 0.735000 | 1.0000 |
| KL20 | 0.003210 | 0.0706 |
| KL21 | 0.523000 | 0.8850 |
| KL23 | 0.255000 | 0.8280 |
| KL24 | 0.539000 | 0.8890 |
| KL25 | 0.177000 | 0.6870 |
| KL27 | 0.085500 | 0.6680 |
| KL3 | 0.150000 | 0.6680 |
| KL30 | 0.772000 | 1.0000 |
| KL39 | 0.652000 | 0.9780 |
| **KL43** | **0.000171** | **0.0113** |
| KL45 | 0.150000 | 0.6680 |
| KL48 | 0.652000 | 0.9780 |
| KL5 | 0.389000 | 0.8280 |
| KL51 | 0.150000 | 0.6680 |
| KL52 | 1.000000 | 1.0000 |
| KL53 | 0.389000 | 0.8280 |
| KL54 | 0.523000 | 0.8850 |
| KL55 | 0.150000 | 0.6680 |
| KL57 | 1.000000 | 1.0000 |
| KL59 | 1.000000 | 1.0000 |
| **KL62** | **0.001140** | **0.0376** |
| KL63 | 1.000000 | 1.0000 |
| KL64 | 1.000000 | 1.0000 |
| KL67 | 0.523000 | 0.8850 |
| KL74 | 1.000000 | 1.0000 |
| KL8 | 0.523000 | 0.8850 |
| KL9 | 1.000000 | 1.0000 |
| Unknown | 0.068300 | 0.6440 |

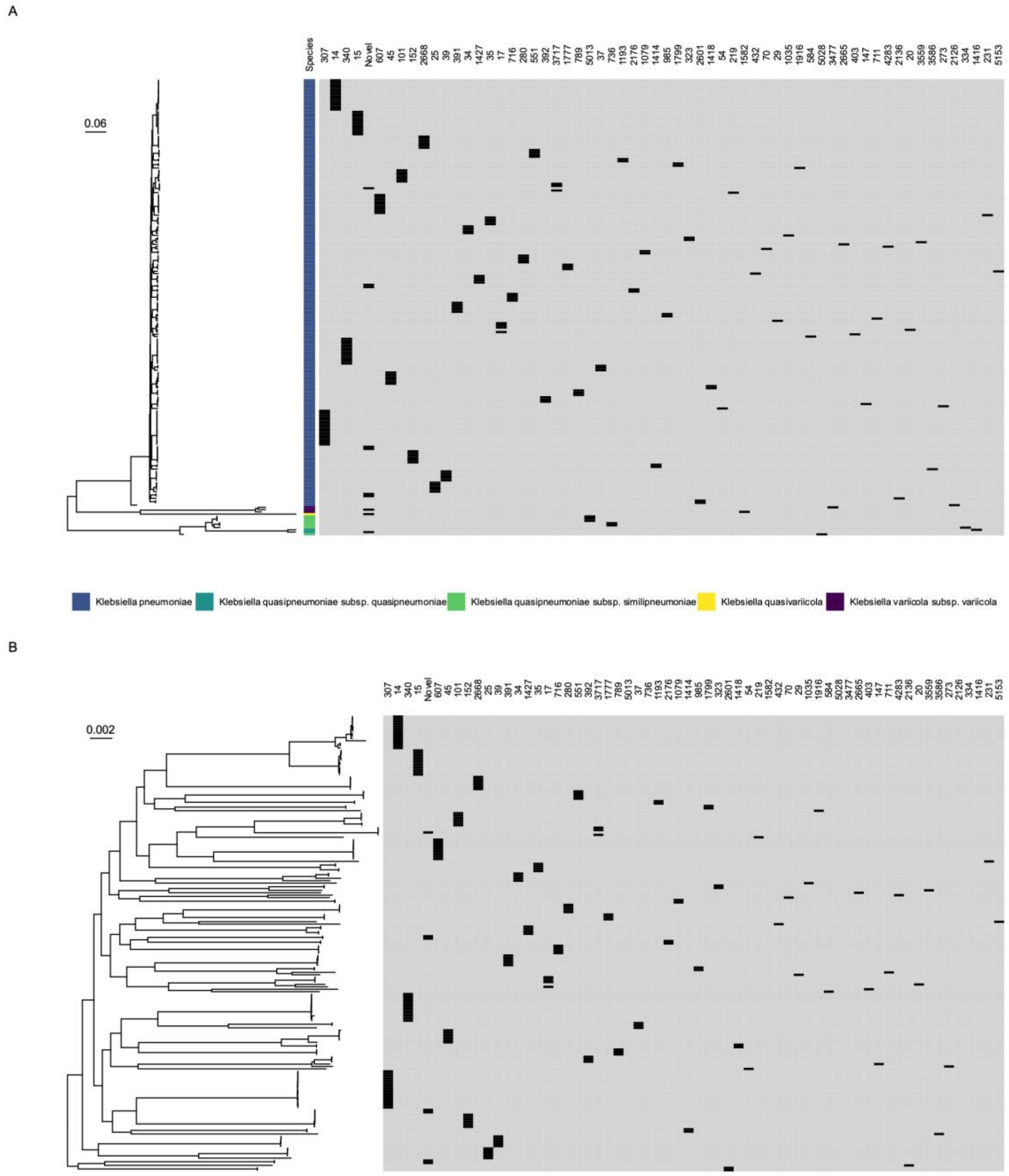


**Supplementary Figure 1:** Midpoint-rooted core-gene maximum-likelihood phylogenetic trees of isolates from the study. (A) shows the full phylogeny with species (coloured bar) and sequence type (ST, black bars); (B) restricts to *K. pneumoniae sensu stricto*, showing STs (black bars). Scale bars show nucleotide substitutions per site.

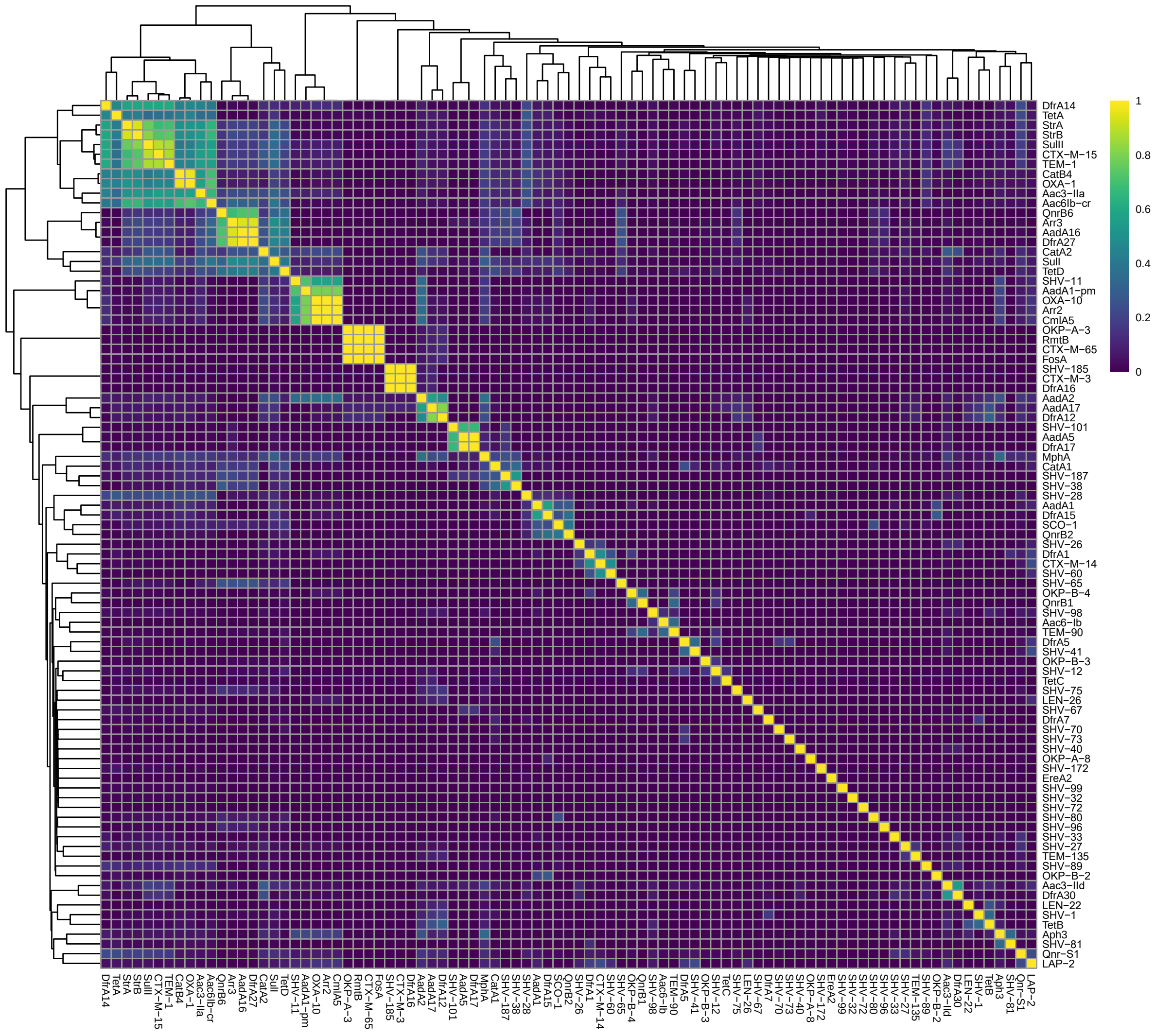


**Supplementary Figure 2:** Jaccard-distance heatmap of presence of ARIBA-identified AMR genes, clustered with a hierarchical clustering algorithm. Several AMR-gene clusters are apparent.

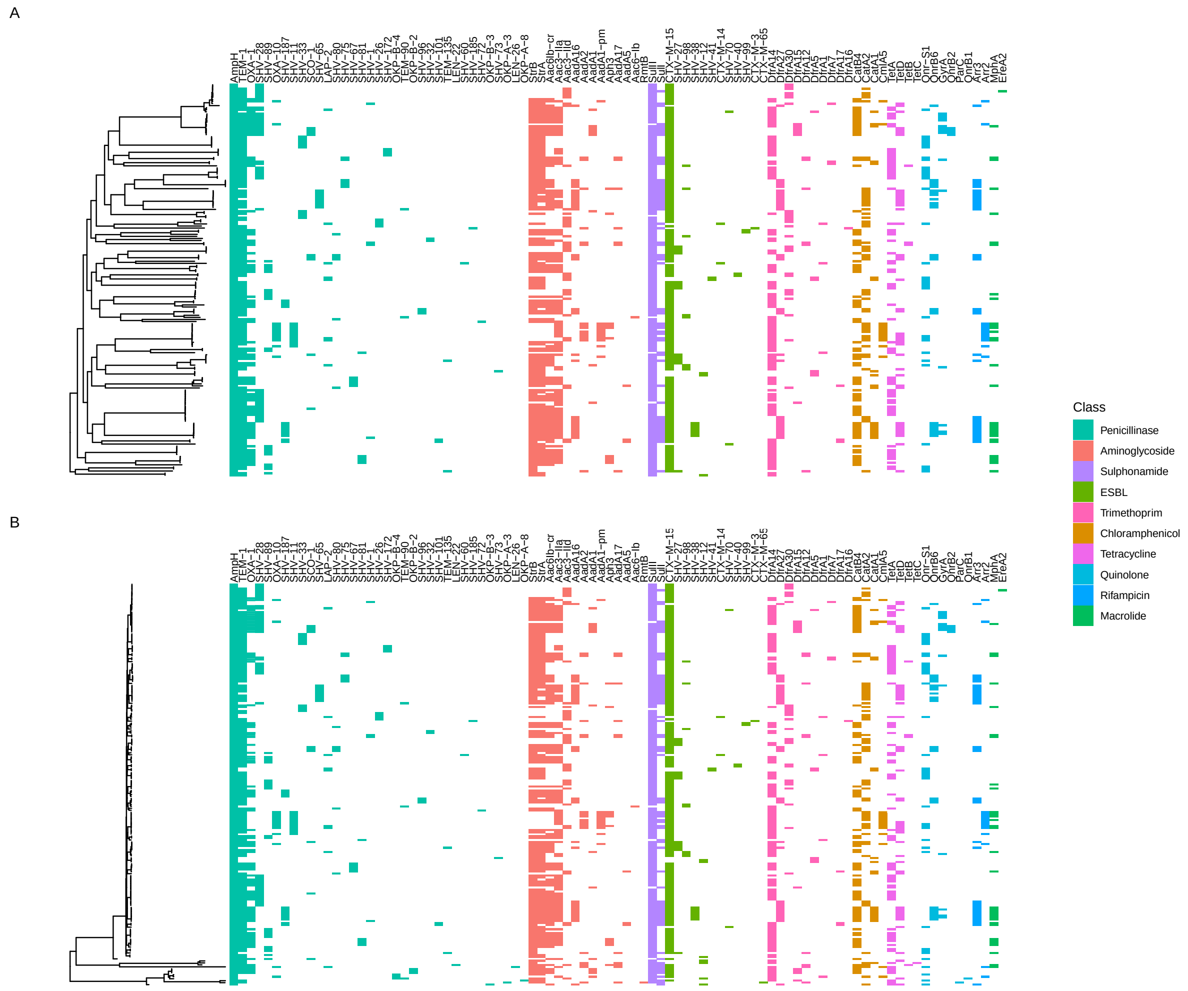


**Supplementary Figure 3:** Presence of ARIBA-identified AMR genes mapped back to phylogeny for (A) KPI isolates only (B) all samples. Some lineage association of AMR genes is apparent.

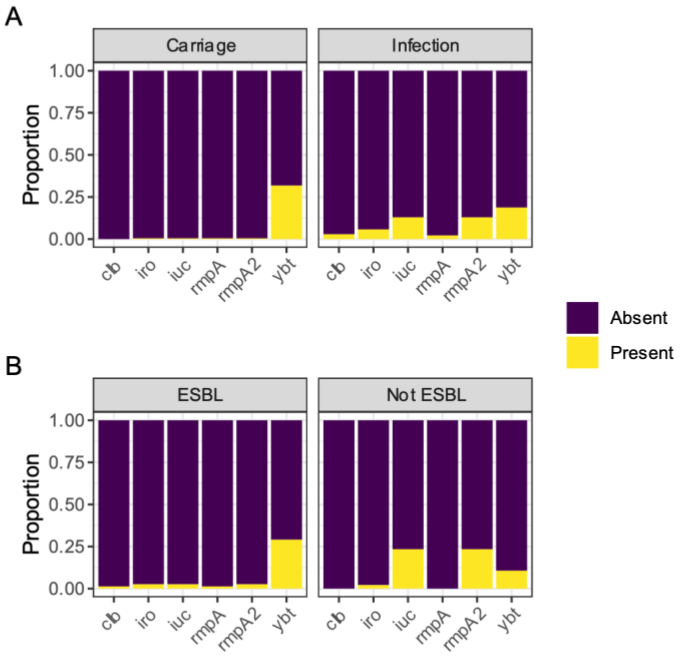


**Supplementary Figure 4:** Virulence determinants of Malawian isolates stratified by (A) carriage or infecting isolate (B) ESBL or non-ESBL isolate
